# Probing local chromatin dynamics by tracking telomeres

**DOI:** 10.1101/2022.02.15.480529

**Authors:** Rebecca Benelli, Matthias Weiss

**Affiliations:** Experimental Physics I, University of Bayreuth, Universitätsstr. 30, D-95447 Bayreuth, Germany

## Abstract

Chromatin dynamics is key for cell viability and replication. In interphase, chromatin is decondensed, allowing the transcription machinery to access a plethora of DNA loci. Yet, decondensed chromatin occupies almost the entire nucleus, suggesting that DNA molecules can hardly move. Recent reports have even indicated that interphase chromatin behaves like a solid body on mesoscopic scales. To explore the local chromatin dynamics, we have performed single-particle tracking on telomeres under varying conditions. We find that mobile telomeres feature in all conditions a strongly subdiffusive, anti-persistent motion that is consistent with the monomer motion of a Rouse polymer in viscoelastic media. In addition, telomere trajectories show intermittent accumulations in local niches at physiological conditions, suggesting the surrounding chromatin to reorganize on these time scales. Reducing the temperature or exposing cells to osmotic stress resulted in a significant reduction of mobile telomeres and the number of visited niches. Altogether, our data indicate a vivid local chromatin dynamics, akin to a semi-dilute polymer solution, unless perturbations enforce a more rigid state of chromatin.

**Statement of Significance:** In interphase cells, chromatin is decondensed and occupies almost the entire nucleus, suggesting DNA molecules to be fairly immobile. Recent reports even indicated chromatin to behave like a solid body on mesoscopic scales. Tracking individual telomeres, we have explored local chromatin dynamics under varying conditions. Our data reveal that telomeres show an antipersistent subdiffusion (fractional Brownian motion) at physiological conditions and at lower temperatures, indicating chromatin to be a viscoelastic fluid on submicron length scales. Telomeres also appear to visit local niches, supposedly provided by the constantly reorganizing chromatin. Applying osmotic stress significantly reduced the fraction of mobile telomeres and the number of visited niches, indicating that chromatin eventually assumes an elastic, solid-like behavior under these conditions.

## I. INTRODUCTION

The nucleus is the defining organelle of eukaryotic cells and responsible for hosting a species-specific number of DNA molecules [1]. Between cell divisions, i.e. in interphase, the DNA appears decondensed with a local organization into (groups of) nucleosomes that is mediated by histones [1]. Nucleosomes have a diameter of 10 nm but contain about 70 nm of DNA, hence facilitating a more compact assembly of DNA than predicted for a simple random coil polymer. As a result, even 2 m of DNA fit into human nuclei with a diameter of few *μ*m while most of the genetic information remains accessible for vital transcription processes. Remarkably, even the decondensed chromatin, filling up almost the complete nucleus, remains dissected into distinct territories [2] that eventually condense into individual chromosomes at the onset of cell division.

The ambivalent nature of interphase chromatin, i.e. its similarity to dynamically reorganizing polymers while maintaining stationary territories, has fueled several studies on elucidating whether chromatin shall be seen as a fluid or rather as a solid [3]. For example, an image correlation analysis has revealed that chromatin regions beyond the boundaries of single-chromosome territories move in a coherent fashion [4], suggesting an elastic coupling between different chromatin loci. Furthermore, a recent study [5], based on fluorescence recovery after photobleaching experiments, has put forward evidence that chromatin has properties of a solid, *in vivo* and *in vitro*. In contrast, an earlier study had supported the picture of liquid chromatin that can even undergo a fluid-fluid phase transition under physiological conditions [6]. In all of these studies, however, microscopy techniques were used that assessed chromatin dynamics only on intermediate to large length scales above the diffraction limit, hence leaving aspects on the submicron scale masked.

Here, we have used single-particle tracking on telomeres in living mammalian cells under several conditions to fill that gap. Telomeres are vital nucleotide sequences at both ends of each chromosome that are involved in a variety of physiological functions [7]. Their motion has been used already as a probe for studying chromatin dynamics *in vivo* [8, 9] and single-particle tracking (SPT) experiments on telomere-bound TRF-2 proteins have revealed that telomeres exhibit a distinct anomalous diffusion characteristics in mammalian cells [9, 10]. In particular, the telomeres’ mean square displacement (MSD) was seen to display a sublinear power-law increase, consistent with the motion of a monomer in a Rouse polymer and partially driven by active noise mediated by the cytoskeleton. Going beyond these results, we show here that the subdiffusion of telomeres bears all features of a fractional Brownian motion (FBM) process with intermittent jumps between apparent niches, supposedly formed by the surrounding chromatin. Upon lowering the temperature or after applying osmotic stress, the diffusion anomaly is further enhanced, jumps are suppressed, and an increasing fraction of immobile telomeres is observed. We conclude from our data that chromatin appears like a viscoelastic fluid on small length scales. At lowered temperatures or after messing up the intracellular water balance, chromatin appears to stiffen and behaves more and more like a solid.

## II. MATERIALS AND METHODS

### A. Cell culture and microscopy

Sample preparation and microscopy were done in accordance with earlier work [10]. In brief, bone osteosarcoma cells (U2OS, DSMZ Cat# ACC-785, RRID:CVCL 0042) were grown at 37^°^C and 5% CO_2_ in McCoy’s medium with phenol red (Biochrom, Germany) supplemented with 10% fetal calf serum (FCS), 1% L-glutamine, 1% sodium pyruvate and 1% penecillin/streptomycin. In accordance with earlier studies [9–11], telomeres were highlighted by a GFP-tagged TRF-2 construct (a kind gift of Y. Garini, Bar-Ilan University, Israel) that recognizes the human telomeric sequences TTAGGG. Transient transfection with the plasmid was performed 24 h prior to microscopy, using Lipofectamine3000 (Thermo Fisher Scientific, Germany) according to the manufactures protocol (4.5 *μ*l Lipofectamine3000 Reagent and 75 *μ*l Opti-MEM mixed with 1 *μ*l P3000 Reagent, 0.5 *μ*g DNA and 75 *μ*l Opti-MEM).

For live-cell microscopy, cells were plated 24 h prior to transfection in 4-well *μ*-slide microscopy chambers (ibidi, Martinsried, Germany) at a density of 50,000 cells/well. For standard imaging, the medium was changed to MEM without phenol red supplemented with 5% FCS and 5% HEPES (referred to as ‘imaging medium’ in the remainder). For hypoosmotic conditions, the medium was changed to 5% McCoy’s medium with 95% ultrapure water (Invitrogen, Waltham, Massachusetts, United States). For hyperosmotic conditions, a solution of 1 M saccharose (Roth, Germany) in McCoy’s medium was prepared and diluted with image medium to the final concentration. The medium was applied 15 min prior to imaging in all cases.

Imaging was performed with a customized spinning-disk confocal microscope based on a DMI 4000 stand (Leica Microsystems, Germany), a CSU-X1 spinning-disk unit (Yokogawa Microsystems, Japan), an HC PL APO 100x/1.40NA oil immersion objective, and an Evolve 512 EMCCD camera (Photometrics, USA). Live-cell imaging was performed at varying temperatures using a custom-made incubation chamber. If the temperature for microscopy deviated from the standard incubation temperature of 37^°^C, cells were placed in the incubation chamber for at least 15 min before imaging started to allow for an adaption. Samples were illuminated at 488 nm and fluorescence light was detected in the range 500-550 nm. Confocal images (single optical slices of the cell) were taken at an interval of Δ*t* = 110 ms and an exposure time of 50 ms for a total time of up to 20 min.

### B. Data processing and analysis

Images were cropped to speed up all subsequent evaluation steps. Telomeres were tracked with a custom-written Matlab script as described before [10]. In particular, tracks were not allowed to include gaps and the estimated step size was chosen ∼ 0.7 *μ*m, i.e. in the range of the detected telomere steps. For sufficient statistics in the analyses, only tracks with at least 1000 successive positions were retained and immobile tracks within this set were discarded (see below). Correcting trajectories for the center-of-mass-movement of the entire nucleus was done as described before [10]. Nuclei that showed a marked rotation over time or suffered from a pronounced drift were not taken into consideration.

Standard measures like the mean square displacement (MSD), the power-spectral density (PSD) etc. were obtained by customized Matlab routines in a straightforward fashion. Analyses of MSDs were performed on full-length trajectories; using only the first 1000 positions did not significantly alter the results (table in the Supplementary Information). To facilitate an ensemble-averaging, correlation functions and PSDs were determined by trimming all trajectories to 1000 positions, i.e. by cutting away additional positions.

The number of blobs in each full-length trajectory, *n*_*B*_, i.e. the number of its apparently separable trajectory segments, was determined by rating local point position accumulations with a customized Matlab routine. First, the mean density of points, *ρ*_0_ = *N/A*_0_, and the associated typical length scale 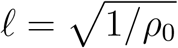, were defined from the number of positions in the trajectory, *N*, and the polygonal area *A*_0_ enclosed by the boundary of all positions. Then, we determined the local point density *ρ*_*i*_ within a circle of radius 5*𝓁*, centered at trajectory positions **r**_*i*_ (1 ≤ *i* ≤ *N*). To avoid errors at the boundary, the intersection of the circle area and *A*_0_ was evaluated if the circle was not completely included in the trajectory’s boundary. All trajectory positions were subsequently assigned coordinates on a square lattice with spacing *𝓁/*2 and all lattice sites were initially assigned a value zero. Lattice sites to which at least one trajectory position **r**_*i*_ with *ρ*_*i*_*/ρ*_0_ *>* 0.9 had been assigned, were set to a value of unity to highlight loci of increased point density. The resulting square image, i.e. the lattice, was subjected to a morphological opening with a 3 × 3 matrix with unity entries on all positions. In a final step, the number of disjoint clusters of unity value in the image were extracted by the Matlab command *bwconncomp*, similar to a Hoshen-Kopelman algorithm evaluation of spin lattices [12]. Examples for this approach are shown in Fig. 3a.

Random walk trajectories of the FBM type, for comparison with experimental data, were obtained with Matlab as described earlier [13].

## III. RESULTS AND DISCUSSION

### A. Telomeres show Rouse dynamics in untreated cells indicating fluid chromatin

To gain insights into the local dynamics of chromatin, we tracked telomeres in U2OS cells (see Materials and Methods for details); representative cell images are shown in the Supplementary Information (Fig. S1). For the analysis, we calculated in the first instance the time-averaged mean square displacement (TA-MSD) for each telomere trajectory as a function of the lag time *τ* = *k*Δ*t*,

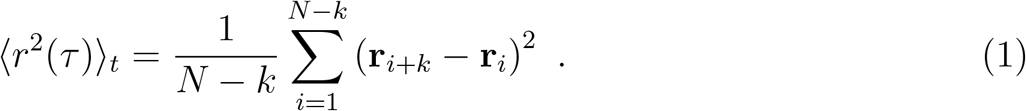

Here, *N* denotes the total length of the individual trajectory unless trajectories are explicitly stated to have been trimmed to a length *N* = 1000.

Virtually all TA-MSDs obtained from cells at physiological conditions featured a clear sublinear characteristics of the TA-MSD on time scales below a few seconds, with considerable variations between individual trajectories (see Fig. 1a). For large lag times, a crossover to steeper slopes is observed, in agreement with earlier observations [9, 10].

**FIG. 1:**
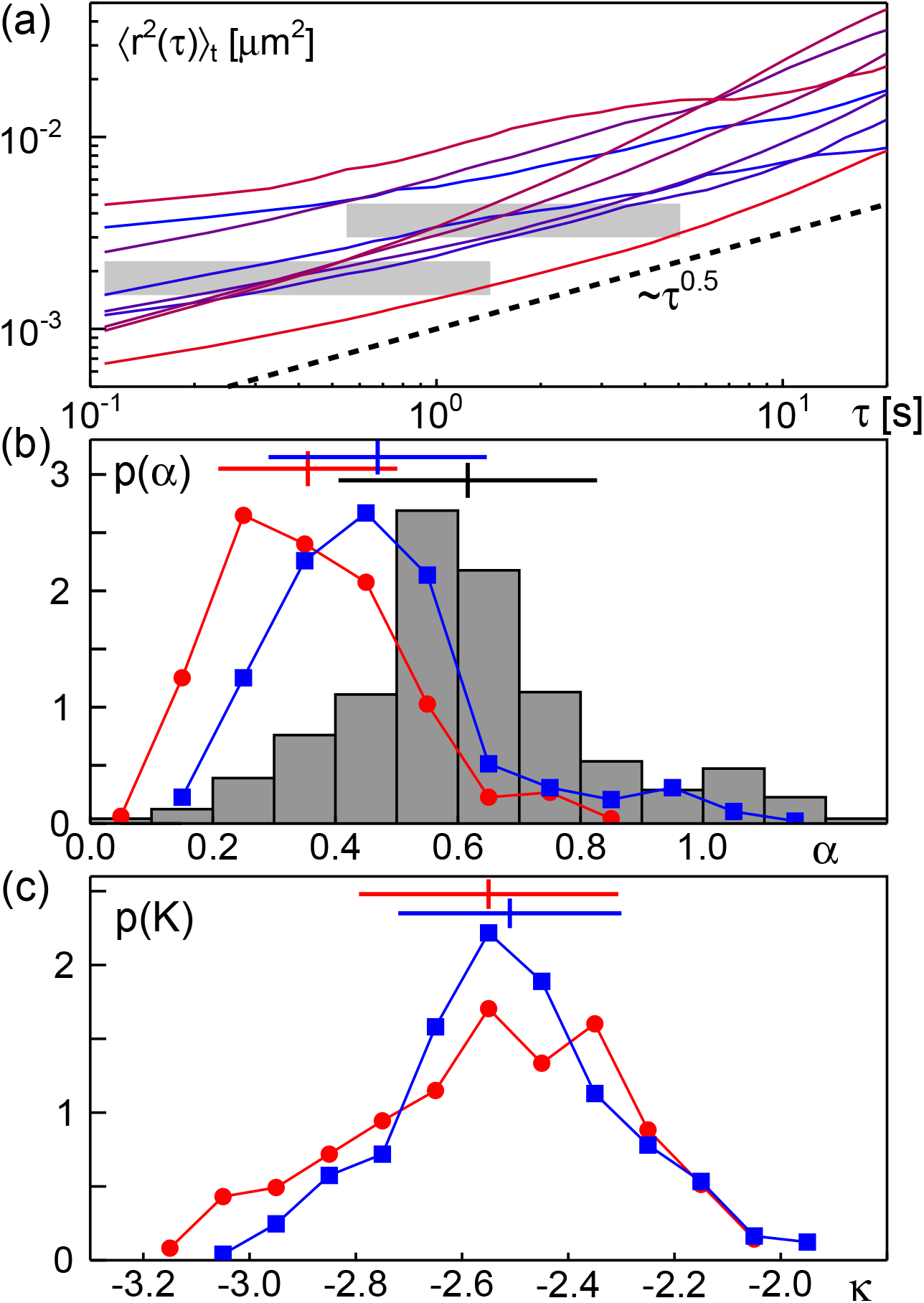
(a) Representative TA-MSDs of telomeres obtained from U2OS cells at physiological conditions. As a guide to the eye, a sublinear power law is shown as dashed line. Fit regions (A) and (B), defined as 1 ≤ *τ/*Δ*t* ≤ 13 and 5 ≤ *τ/*Δ*t* ≤ 46, are highlighted by horizontal grey bars. (b) The PDF of scaling exponents, *p*(*α*), as obtained by fitting TA-MSDs in the intervals (A) and (B) (red circles and blue squares) feature a similar width *σ* but different means ⟨*α*⟩ (indicated by haircrosses). In line with the anticipated effect of a static localization error, a larger mean is obtained for fit interval (B). Using a resampling approach yields a considerably wider PDF (grey histogram) with an even more elevated mean. See also main text for discussion. (c) The associated PDF of generalized diffusion coefficients, *p*(*K*), for convenience shown here as a function of *κ* [Eq. (3)], features a roughly lognormal shape with a similar mean for both fitting intervals.

To obtain a quantitative and objective measure for the variability of TA-MSDs on short time scales, we have performed a linear regression of log(⟨*r*^2^(*τ*)⟩_*t*_) versus log(*τ*) for each trajectory, hence assuming a power-law scaling

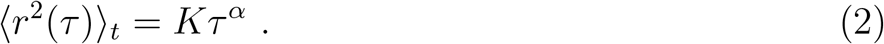

Here, *K* is a generalized diffusion coefficient with units of an area per fractional time, determined via the scaling exponent 0 *< α* ≤ 2. Standard Brownian motion is recovered for *α* = 1, ballistic motion yields *α* = 2.

Extracting the scaling exponent by applying this fitting procedure to the interval 5Δ*t* ≤ *τ* ≤ 38Δ*t*, we classified trajectories with *α <* 0.1 as immobile and disregarded them for subsequent analyses. A similar approach had been taken in Ref. [14]. The fraction of mobile trajectories, *f*_mobile_, was 100% for cells at physiological conditions (corresponding to several hundred trajectories), but varied for other conditions (Table I).

**TABLE I:** Mean anomaly exponents ⟨*α*⟩ and logarithmized diffusion coefficients ⟨*κ*⟩ [Eq. (3)] with the respective standard deviation *σ* of the associated PDF at the indicated conditions as determined in the fit ranges (A) and (B), defined in the main text. The fraction of trajectories that have been classified as mobile for this analysis is given as *f*_mobile_ with the corresponding number of full-length trajectories stated below in brackets.

| condition | $f_{\text{mobile}}$ | fit range | $\langle\alpha\rangle \pm \sigma$ | $\langle\kappa\rangle \pm \sigma$ |
| --- | --- | --- | --- | --- |
| 37°C | 100% | (A) | $0.36 \pm 0.15$ | $-2.55 \pm 0.24$ |
| | (487) | (B) | $0.47 \pm 0.18$ | $-2.51 \pm 0.21$ |
| 34°C | 96.4% | (A) | $0.33 \pm 0.16$ | $-2.32 \pm 0.32$ |
| | (1052) | (B) | $0.41 \pm 0.17$ | $-2.51 \pm 0.26$ |
| 30°C | 97.5% | (A) | $0.28 \pm 0.12$ | $-2.49 \pm 0.26$ |
| | (1264) | (B) | $0.35 \pm 0.11$ | $-2.61 \pm 0.28$ |
| 25°C | 88.4% | (A) | $0.28 \pm 0.13$ | $-2.57 \pm 0.29$ |
| | (831) | (B) | $0.36 \pm 0.13$ | $-2.68 \pm 0.23$ |
| hyper.<br>(0.25 M) | 61.7% | (A) | $0.22 \pm 0.08$ | $-2.17 \pm 0.28$ |
| | (385) | (B) | $0.22 \pm 0.10$ | $-2.39 \pm 0.48$ |
| hyper.<br>(0.5 M) | 16.6% | (A) | $0.14 \pm 0.07$ | $-2.69 \pm 0.33$ |
| | (166) | (B) | $0.18 \pm 0.08$ | $-2.66 \pm 0.48$ |
| hyper.<br>(1 M) | 14.4% | (A) | $0.09 \pm 0.06$ | $-2.63 \pm 0.32$ |
| | (157) | (B) | $0.13 \pm 0.10$ | $-2.67 \pm 0.27$ |
| hypo. | 89.9% | (A) | $0.10 \pm 0.06$ | $-2.81 \pm 0.40$ |
| | (310) | (B) | $0.14 \pm 0.09$ | $-2.97 \pm 0.29$ |

The pool of mobile trajectories was then subjected to a deeper TA-MSD analysis via the same fitting approach, using two different lag time intervals, named (A) and (B) hereafter, that allow for a first assessment whether MSDs are influenced by localization errors. In particular, we used for each trajectory the intervals (A) Δ*t* ≤ *τ* ≤ 13Δ*t* and (B) 5Δ*t* ≤ *τ* ≤ 46Δ*t* for retrieving *α* and *K* (cf. also Fig. 1a).

The obtained probability density function (PDF) of anomaly exponents, *p*(*α*), shows a pronounced peak well below unity for both fitting ranges, confirming the overall sublinear growth of TA-MSDs, i.e. a subdiffusive characteristics, on short and intermediate time scales. Beyond this qualitative agreement, fitting ranges (A) and (B) yielded markedly different means, ⟨*α*⟩, but a similar width (i.e. standard deviation *σ*) of the PDF (Fig. 1b and Table I). In particular, a mean exponent ⟨*α*⟩ ≈ 0.36 was obtained for fitting range (A) whereas fitting in interval (B) yielded ⟨*α*⟩ ≈ 0.47, the latter of which is in good agreement with earlier observations [9, 10].

The PDF of generalized diffusion coefficients, *p*(*K*), obtained from fitting in intervals (A) or (B) both feature a roughly lognormal shape (see [15] for a discussion of this fairly common observation) with comparable peak positions but slightly different widths (Fig. 2c). Please note: Due to the lognormally varying values for the generalized diffusion coefficient *K* and the dependence of its units on the scaling exponent *α*, we have chosen to use the dimensionless quantity

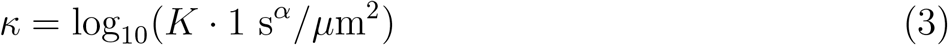

for quantitative comparisons (cf. Fig. 1c and Table I).

**FIG. 2:**
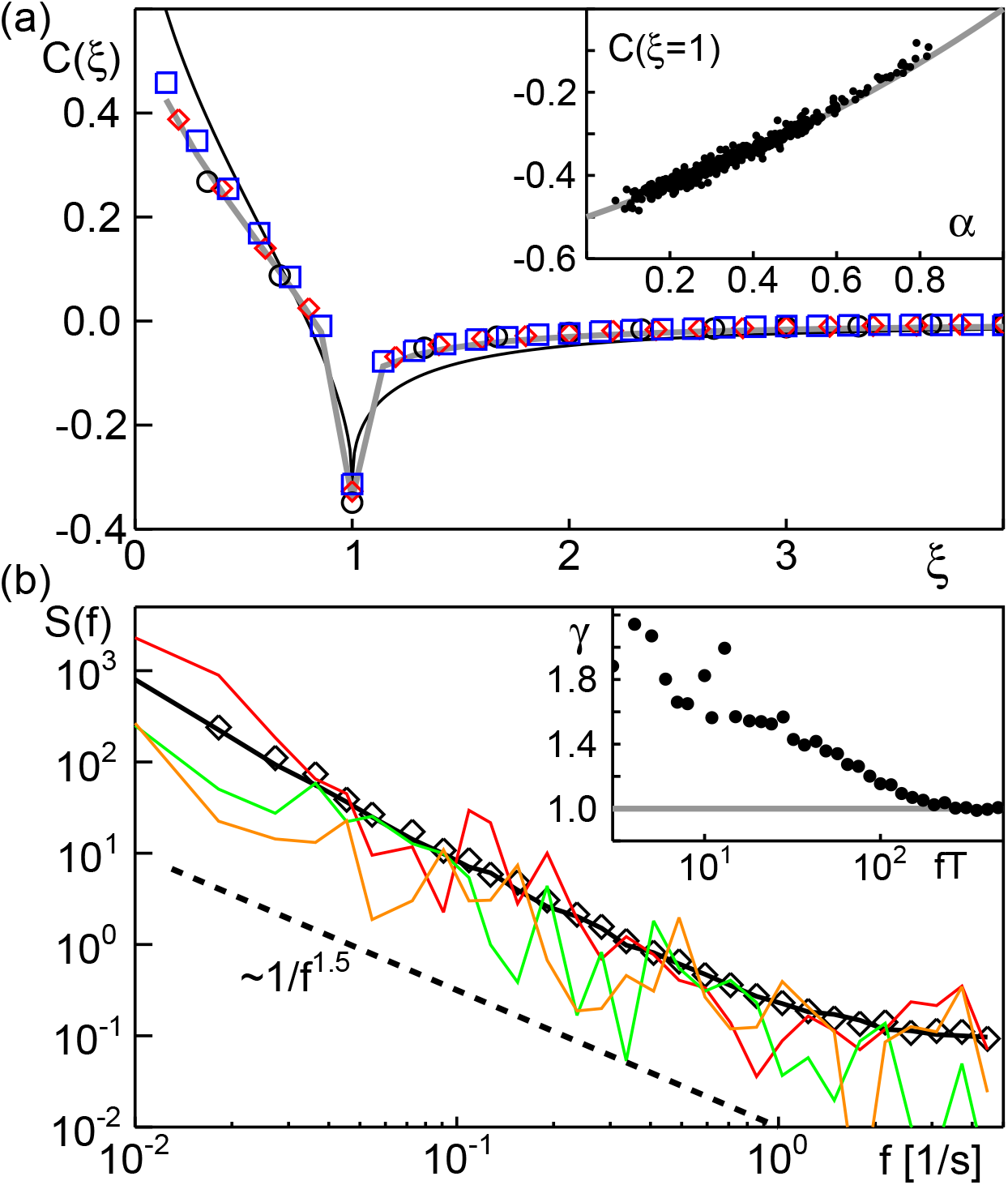
(a) The ensemble- and time-averaged VACF [Eq. (5)] of telomeres in untreated cells at 37°C, shown for *n* = 3, 5, 7 (circles, diamonds, squares), is well captured by the analytical prediction for FBM in the presence of a static localization error (grey line). Deviations to the prediction of an unperturbed FBM random walk (thin black line) are clearly visible. Inset: The value *C*(*ξ* = 1) for *n* = 3 changes with the anomaly exponent *α* (from fit interval (B)) as predicted for FBM processes (grey line). (b) Ensemble-averaged PSDs for *x*- and *y*-coordinates (black line and diamonds) follow the anticipated power-law decay (dashed line), about which the PSD of individual trajectory coordinates fluctuate (examples shown in different colors). Inset: The coefficient of variation *γ* converges to unity for large frequencies, as predicted for FBM processes.

The observation of an increasing ⟨*α*⟩ for fit ranges with larger lag times strongly indicates that trajectories are plagued by a remnant static localization error, i.e. particle positions are retrieved from images with a non-negligible random error (see [13, 16] for a discussion on the influence of localization errors in subdiffusive TA-MSDs). As a consequence, TAMSDs will converge to a positive constant for *τ* → 0 and fitting with a simple power law [Eq. (2)] results in an apparently lower value of the scaling exponent. When fitting on larger time scales, the localization error successively becomes negligible, resulting in larger values of *α*, in accordance with our observations (Fig. 1b). Applying a resampling approach [13], supposed to erradicate the influence of localization errors, even yielded a PDF with a mean ⟨*α*⟩ ≈ 0.62 (Fig. 1b). While this result supports the assessment that trajectories include a non-vanishing static localization error, the considerably wider PDF obtained with this approach suggests that the mean might be less accurate. Given this caveat and since the resampling approach does not retrieve the generalized diffusion coefficient, *K*, we have used the results obtained from fit interval (B) as benchmark for all subsequent comparisons and discussion.

We would like to emphasize at this point that very similar results for the PDFs of *α* and *K* were obtained when trimming all trajectories to the same length (*N* = 1000) by cutting away any exceeding positions from full-length trajectories (see Table in the Supplementary Information). In particular, we did not observe a significant dependence of *α* on the trajectory length.

Taken together, our data obtained for cells at physiological conditions are in good agreement with earlier results that have revealed that telomeres show a distinct subdiffusion with an average scaling exponent near to *α* = 1*/*2, as predicted for the motion of a monomer in a Rouse polymer [9, 10]. Taking this interpretation further, telomeres should show an anti-persistent random walk with all properties of fractional Brownian motion (FBM) for lag times lower than the Rouse time, i.e. successive steps should not be chosen in a Markovian fashion but rather with an anti-persistent memory kernel.

To probe the existence of such an anti-persistent memory kernel, we considered the time-averaged velocity autocorrelation function (VACF)

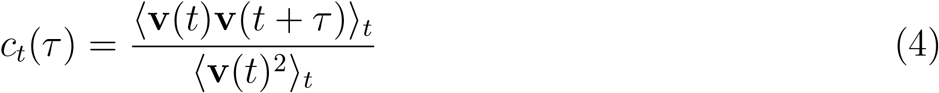

for each trajectory as well as the considerably smoother ensemble- and time-averaged VACF

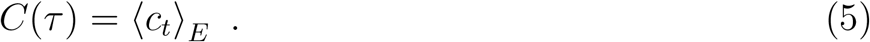

Here, *τ* = *k*Δ*t* is again the lag time while **v**(*t*) = [**r**(*t*+*δt*)−**r**(*t*)]*/δt* denotes the instantaneous velocity, based on the spatial increment **r**(*t* + *δt*) − **r**(*t*) taken within an integer multiple of the frame time, *δt* = *n*Δ*t*. To allow for an easy averaging, we have trimmed all trajectories to a common length *N* = 1000 for these quantities.

For a comparison to FBM, it is convenient to express the VACF in terms of a rescaled lag time *ξ* = *τ/δt* = *k/n* since antipersistent random walks of the FBM type feature a distinct minimum *C*(*ξ* = 1) = 2^*α*−1^ − 1 [17, 18]. Indeed, extracting *c*_*t*_(*ξ* = 1) for each telomere trajectory (using *n* = 3 for determining the instantaneous velocity) and plotting it versus the apparent scaling exponent *α*, obtained from the previously defined benchmark interval (B), nicely complies with this prediction (Fig. 2a, inset), providing more evidence that telomeres perform an FBM-like motion.

Moreover, the rather smooth ensemble- and time-averaged VACF [Eq. (5)] confirms both, an FBM-like motion of telomeres and the presence of a non-negligible localization error. While a clear antipersistence dip is observed at *ξ* = 1 for different choices of *δt* = *n*Δ*t* (Fig. 2a), also marked deviations from the analytical prediction for a fully self-similar FBM are seen. In fact, an analytical expression for the VACF of FBM random walks that explicitly accounts for localization errors has been derived for *δt* = *n*Δ*t* and lag time *τ* = *k*Δ*t* [16]:

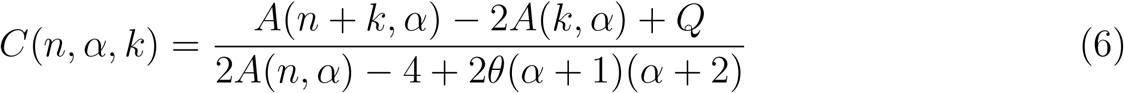

with

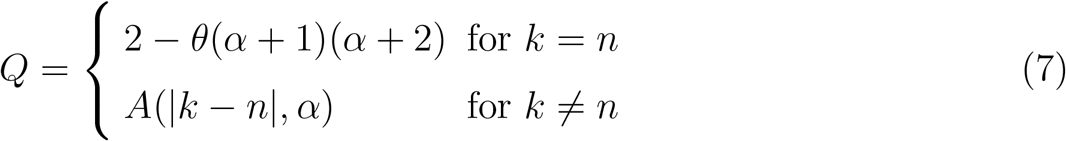

and

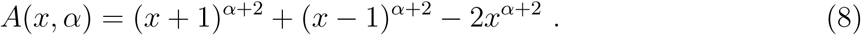

Here, the constant 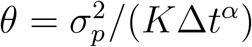 summarizes the influence of the static localization error (i.e. the variance 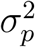 of the random scattering around the true position), while a dynamic localization error is included via sums of the functions *A*(*x, α*). For nonzero values of *θ*, the exact self-similarity of FBM random walks is broken, resulting in a reduction of the anti-persistent dip in the VACF for increasing *n*. This function almost perfectly describes the experimental data (Fig. 2a), confirming our telomere trajectories at physiological conditions to be FBM-like with a static localization error. We would like to highlight at this point that including a mild static localization error does not invalidate the aformentioned relation *C*(*ξ* = 1) = 2^*α*−1^ − 1 as the perturbations of *α* and *C*(*ξ* = 1) approximately cancel out (see [19] for details).

To obtain a last piece of evidence that telomeres move according to an FBM process, we have determined the power-spectral density (PSD) of the *x*- and *y*-coordinates of all trajectories after trimming them to a common length *N* = 1000 to simplify the analysis. To account for locus-dependent variations of the generalized diffusion coefficient, each coordinate of each trajectory was rescaled by the root-mean-square step length between successive frames. The resulting ensemble-averaged PSDs for each spatial coordinate were almost identical and followed the anticipated power-law decay *S*(*f*) ∼ 1*/f* ^*β*^ with *β* = *α* + 1 [20]. PSDs of individual trajectories fluctuated randomly around these ensemble averages (see Fig. 2b), as expected. These fluctuations can be quantified by the coefficient of variation, *γ*(*f*) = *σ/μ*, defined as the ratio of the standard deviation *σ*(*f*) of trajectory-wise PSDs at each frequency and their ensemble average, *μ*(*f*).

Plotting *γ* versus the dimensionless frequency *fT* = 1, 2, …, *N*, subdiffusive FBM random walks have been shown to eventually converge to *γ* = 1. Indeed, the data for telomeres follows this very prediction (Fig. 2b, inset), hence strengthening the claim that telomeres move according to an FBM process at physiological conditions.

We next sought to gain information on larger length scales. A visual inspection suggested that trajectories with much more than 1000 positions appear to consist of different blobs (see Fig. 3a for an example), as if telomeres visited different local niches that are provided by the surrounding chromatin. To quantify this, we developed an image-analysis approach that mimics the visual assessment, i.e. accumulations of positions separated by sparsely populated areas are identified as individual blobs (cf. Materials and Methods and examples in Fig. 3a). Certainly, this approach of identifing blobs requires, like any statistical method, the definition of thresholds. Therefore, the identified number of blobs within a trajectory might change in dependence of this parameter. Yet, as long as all trajectories are rated with the same meaningful threshold (selected here to be in line with a visual assessment), the quantification allows for a reasonable comparison between trajectories obtained from different conditions.

**FIG. 3:**
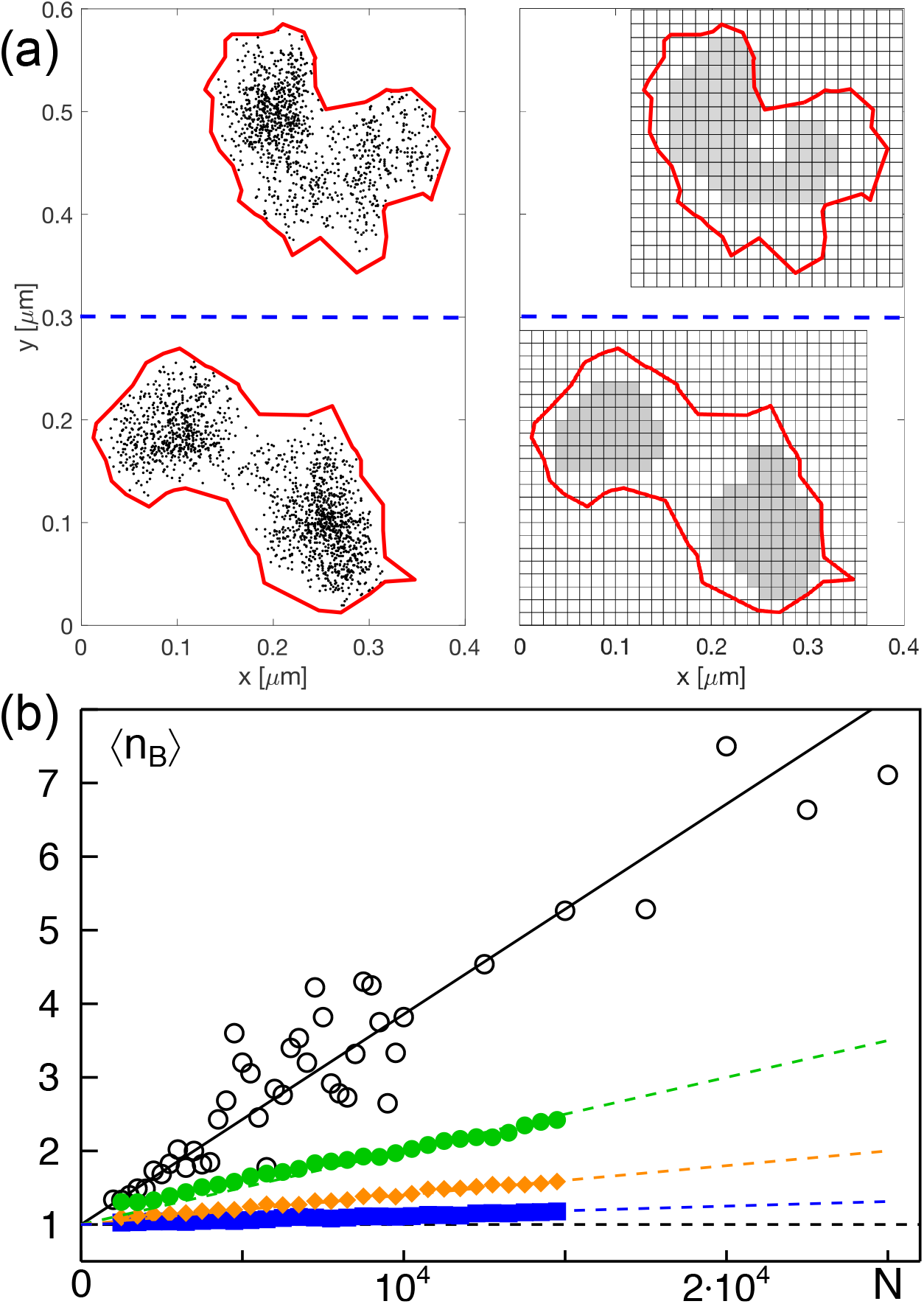
(a) Two representative trajectories (clouds of black dots, with their respective envelope indicated in red) are classified by our automatic image analysis to consist of one and two blobs, respectively (see grey-shaded regions on the right). (b) The average number of blobs ⟨*n*_*B*_⟩ within a trajectory shows a roughly linear increase with the trajectory length *N* in cells at physiological conditions (open black circles and full black line). For very long trajectories, considerable fluctuations around this trend are observed. Simulated FBM trajectories with *α* = 0.4, 0.5, 0.6 (blue, orange, and green symbols and lines) also show a linear increase, yet with a significantly lower slope.

Dissecting full-length trajectories from cells at physiological conditions with this approach, we observed that the mean number of detected blobs, ⟨*n*_*B*_⟩, grows approximately linearly with the trajectory length *N*, albeit with considerable fluctuations for large *N* (Fig. 3b). The latter also includes the successively poor number of trajectories with large *N*. In contrast, ensembles of simulated FBM random walks with *α* = 0.4, 0.5, 0.6 show a considerably lower number of blobs at the same trajectory lengths (Fig. 3b), remaining close to a single blob. The latter would have been the naive guess for a simple random walk. Thus, telomeres on average move according to an FBM but appear to explore different spatial niches that are imprinted by the surrounding, e.g. by neighboring chromatin segments that form transient cages.

Integrating our results up to this point, we conclude that chromatin, as seen via telomere tracking, is not a solid on short and intermediate time and length scales in cells at physiological conditions. It rather bears similarities to a semi-dilute polymer solution for which the Rouse model is an adequate description. Therefore, chromatin appears fluid on local scales with a viscoelastic characteristics on larger scales. In fact, our data and this interpretation are in good agreement with a recent study on the local chromatin viscoelasticity [21]. Given that chromosomes do not reach the state of entanglement but rather remain separated in individual territories, it is also straightforward to conceive a formation of transient voids by surrounding polymer segments that can be explored by individual DNA segments, e.g. telomeres. Forcing chromatin into a more compact state, e.g. by lowering the temperature or applying osmostic stress, should lead to significant deviations in the measures shown in this section.

### B. Altered telomere dynamics at lower temperatures indicates viscoelastic chromatin

To gain insights into local chromatin dynamics after perturbation, we have tracked telomeres at lowered temperatures. Based on earlier observations [4, 10], we reasoned that any active process that drives chromatin dynamics will be mediated by ATPases or GTPases, and hence an Arrhenius behavior of the telomeres’ mobility might be observed. Aiming to maintain all vital functions, the accessible range of lower temperatures was limited to a minimum of about 25^°^C.

As a result, we observed for lower temperatures a slight growth of immobile trajectories and a mild decrease of the average scaling exponent ⟨*α*⟩ for the mobile pool of trajectories (Fig. 4a and Table I). The associated generalized diffusion coefficient did not show major changes (Fig. 4a and Table I). Other measures that reveal the FBM-like nature of the telomeres’ motion, derived from VACFs and PSDs, were basically unchanged (see Fig. S2 in Supplementary Information). Therefore, an Arrhenius-like change of *K* could not be detected yet the anti-persistence of the FBM-like motion was preserved with a markedly lower scaling exponent.

**FIG. 4:**
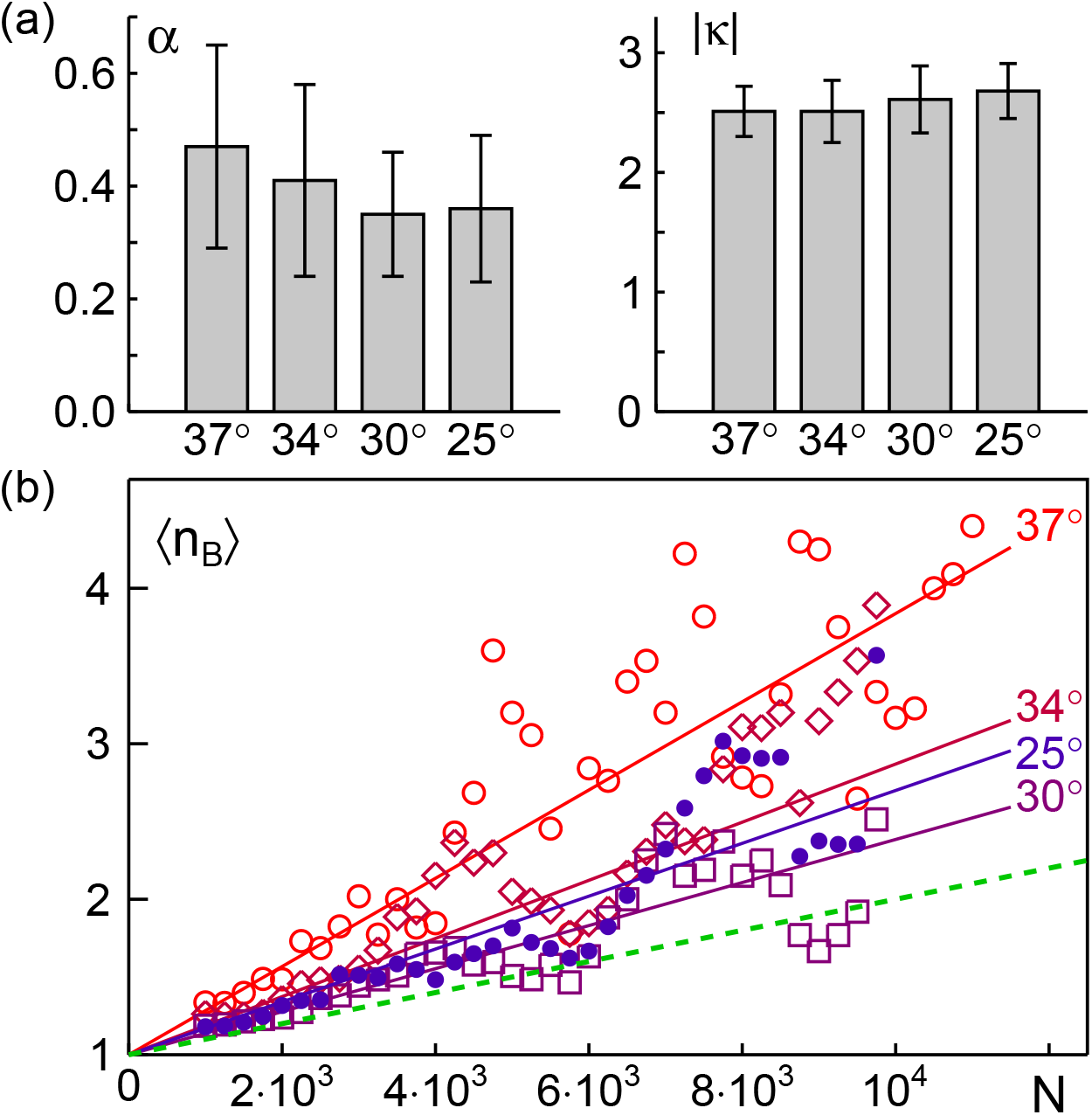
(a) Change of the mean anomaly exponent ⟨*α*⟩ and mean logarithmic diffusion coefficient *κ* [Eq. (3)] at different temperatures. Error bars denote the standard deviation of the associated PDF. Testing any two PDFs *p*(*α*) from different conditions were rated to be significantly different by a Kolmogorov-Smirnov test at a level *p <* 0.001. (b) The mean number of blobs, ⟨*n*_*B*_⟩, as a function of trajectory length, *N*, decreases with temperature but stays above the result for FBM trajectories with *α* = 0.6 (dashed green). Full lines were obtained by linear regression of the respective data sets.

Relating this finding to the above interpretation of chromatin behaving like a semi-dilute polymer solution, the change of ⟨*α*⟩ can be rationalized: Telomeres move like monomers of a Rouse polymer, performing an FBM random walk with an MSD scaling exponent *α*_Rouse_ = 1*/*2 for lag times below the Rouse time scale. The environment for this local motion, however, may not just be purely viscous but the surrounding chromatin may change its conformation at lower temperatures, creating effectively a viscoelastic medium for telomere motion. In viscoelastic media, the complex shear modulus scales as |*G*(*ω*)| ∼ *ω*^*β*^ for shearing frequency *ω*. Therefore, an FBM random walk with an intrinsic anti-persistent memory kernel is additionally equipped with a memory contribution from the surrounding medium [22]. The effective MSD scaling exponent then becomes the product of *β* and the MSD scaling exponent of the pure FBM process [22], leading to *α* = *βα*_Rouse_ *<* 1*/*2 for telomeres. The observed reduction of ⟨*α*⟩ for telomeres at lower temperatures therefore may reflect an increased viscoelasticity of the surrounding, i.e. a lower value of *β*, which suggests that neighboring chromatin regions are less viscous and more elastic at lowered temperatures. In other words, chromatin appears to stiffen locally at lower temperatures but remains a (non-Newtonian) fluid.

This interpretation also predicts that for decreasing temperatures a visiting of different niches should becomes less likely, hence reducing the average number of blobs in trajectories. Indeed, our experimental data support this very expectation. As compared to 37°C, lower temperatures feature smaller average blob numbers ⟨*n*_*B*_⟩ but are still above the result predicted for FBM random walks (Fig. 4b). While the data points and the associated linear fits reveal an effect for temperature changes as low as 3°C, strong fluctuations of ⟨*n*_*B*_⟩ unfortunately do not allow a sufficiently fine assessment of the effect as a function of temperature. Still, the overall reduction of ⟨*n*_*B*_⟩ is clearly visible.

Based on these data, we conclude that chromatin dynamics, as seen via telomere trajectories, maintains its basic FBM-like features for lower temperatures but becomes more anti-persistent and allows for less excursions on the mesoscale. Altogether, this suggests that chromatin as a whole becomes a more elastic and less viscous fluid at lower temperatures.

### C. Telomere dynamics of osmotically stressed cells suggests solid-like chromatin

We next aimed at probing more challenging situations for which a solid-like behavior of chromatin can be expected. In particular, we applied hyper- and hypoosmotic stress to cells (see Materials and Methods) for which changes in nuclear shape and chromatin organization are expected [23, 24]. In fact, diffusion of inert tracers in the nucleus of mammalian culture cells has been shown before to strongly change upon hyperosmotic stress [25] with chromatin assuming a considerably more compact arrangement [26].

As a result of our experiments, we observed for hyperosmotically stressed cells a strong and osmolarity-dependent increase in the number of trajectories that were classified as immobile (Table I). This was accompanied by a significant reduction of ⟨*α*⟩ but only a slight change of the generalized diffusion cofficient for the remaining pool of mobile telemores (Fig. 5a and Table I). Hence, telomeres seem to be mostly arrested in hyperosmotically stressed cells, presumably due to the surrounding chromatin having attained an almost elastic and non-viscous state with a scaling exponent *β* ≪ 1 of the complex shear modulus. In line with this notion, also the average blob number, ⟨*n*_*B*_⟩ was strongly reduced, staying near to unity for all availabe trajectory lengths (Fig. 5b).

**FIG. 5:**
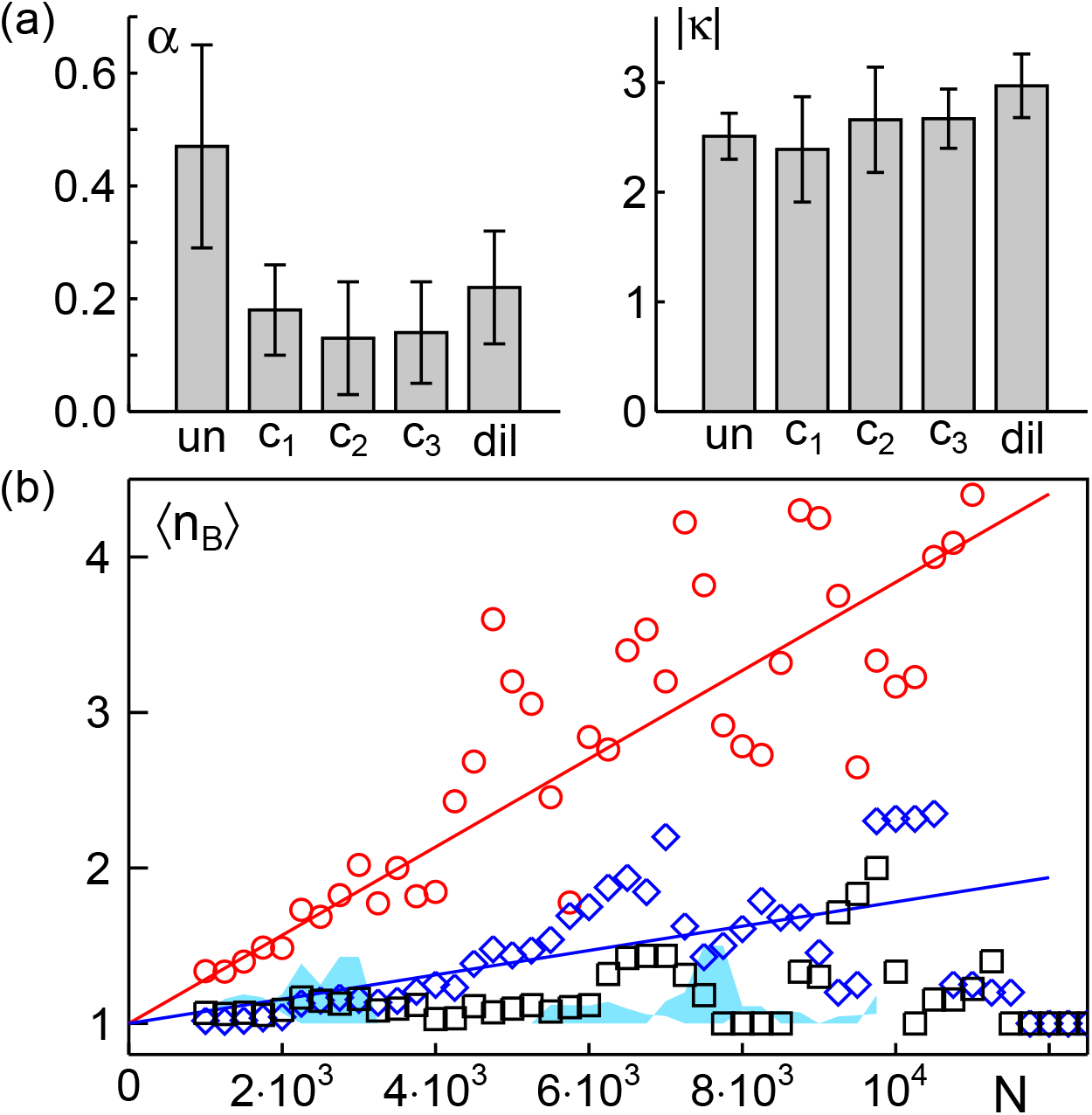
(a) Change of mean anomaly exponent (*α*) and mean logarithmic diffusion coefficient *κ* [Eq. (3)] at different conditions (un: untreated; *c*_1_, *c*_2_, *c*_2_: hyperosmotic shock with 250 mM, 500 mM, or 1 M sugar; dil: hypoosmotic shock). Error bars denote the width of the associated PDFs. Testing any two PDFs *p*(*α*) from different conditions were rated to be significantly different by a Kolmogorov-Smirnov test at a level *p <* 0.001. (b) The mean number of blobs, ⟨*n*_*B*_⟩, observed for different trajectory lengths decreases rapidly upon applying any osmotic shock. A strong reduction as compared to untreated cells (red line) is observable already for condition *c*_1_ (blue line). For larger osmotic shocks (blue background) and dilution (black symbols), virtually no increase in blob number beyond unity can be detected.

Interestingly, applying hyposmotic conditions yielded similar results albeit cells and nuclei clearly showed a swollen phenotype, i.e. the amount of intracellular water was strongly increased. Like in hyperosmotic conditions, ⟨*α*⟩ and ⟨*n*_*B*_⟩ were strongly reduced as compared to untreated cells (Fig. 5 and Table I). This observation suggests that an increased water content in the cytoplasm effectively yields a hyperosmotic stress on chromatin as well, leading to a more elastic and less viscous state in which most of the motion on short and intermediated length and time scales are stalled.

Combining these results, we conclude that a sufficiently strong osmotic stress leads to an apparently more compact and almost rigid chromatin on length scales that are explored by telomeres. In relation to untreated cells, the chromatin appears to be solid-like under these conditions.

## IV. CONCLUSION

Based on our data we conclude that telomeres show an antipersistent subdiffusion of the FBM type at physiological conditions and at lower temperatures, indicating that chromatin has properties of a viscoelastic fluid on submicron length scales. Altogether, chromatin dynamics, as evidenced by the motion of telomeres, appears consistent with the dynamics of polymers under semi-dilute conditions. Applying osmotic stress renders the majority of telomeres immobile and strongly enhances the anti-persistence of the remaining mobile ones, indicating that chromatin assumes an elastic, solid-like behavior on short and intermediate length and time scales under these conditions. Therefore, on length and time scales explored by telomeres, chromatin exhibits features of a viscoelastic fluid at physiological conditions, but stiffens considerably upon osmotic stress.

## Author Contributions

MW conceived the study, RB performed all experiments; RB and MW analyzed the data and wrote the manuscript.

## Acknowledgements

Financial support by the VolkswagenStiftung (Az. 92738) and by the Elite Network of Bavaria (Study Program Biological Physics) are gratefully acknowledged.

